# Cholecalciferol Ameliorates Cyclophosphamide-induced Behavioural Toxicities in Wistar Rats

**DOI:** 10.1101/2025.09.11.675651

**Authors:** Dosunmu Damilare Paul, Ojo Foluso Olamide, Onaolapo Adejoke Yetunde, Onaolapo Olakunle James

## Abstract

Cholecalciferol (Vitamin D_3)_ is a dietary supplement that has been shown to play crucial roles in brain development, immune regulation, neurotransmission and neuroprotection. While there are reports that it influences cognition and mental health there is a dearth of information on its benefits in mitigating cyclophosphamide induced behavioural deficits. This study investigated the possible protective effects of cholecalciferol supplementation on cyclophosphamide induced alterations in open field behaviours, spatial working memory and anxiety-related behaviours in rats. Sixty rats were randomly assigned into six groups (n=10). Groups A and D served as normal and cyclophosphamide control respectively and were fed standard rat chow, groups B and E received Vitamin D_3_ (300 IU/kg), while groups C and F received Vitamin D_3_ (600 IU/kg). Animals in group A-C received intraperitoneal normal saline on day1, while groups D-F got intraperitoneal Cyclophosphamide (100 mg/kg/day on day1). Standard diet and Vitamin D_3_ supplementation were administered daily for 15 days. At the end of the experimental period, animals were exposed to the behavioural paradigm (open field, Y-maze, and elevated plus maze). Vitamin D_3_ mitigated CYP-induced anxiogenic behaviour (dose-dependent). It also significantly improved spatial working memory. The results highlight Vitamin D_3_’s ability to influence CYP-induced loss in cognition in rats via its anxiolytic, cognitive and immune modulatory properties. They also harness Vitamin D_3_’s potential as a possible adjunct in cancer chemotherapy. However, further research will be needed to specify its exact role in cancer chemotherapy.

## 1. Introduction

Increasing prevalence of cancer, a group of diseases marked by the uncontrolled growth and spread of abnormal cells, often resulting in life-threatening conditions has become a global health burden [1]. Current data consistently ranks cancer as the second leading cause of mortality worldwide [2]. Encouragingly, advances in cancer treatment have led to significantly improved patient outcomes and a growing population of cancer survivors [3–5]. However, longer survival is often accompanied by long-term physical, neurological, and psychosocial adverse effects of cancer chemotherapy [6–11]

Of particular concern are deficits in executive functioning, including memory loss and anxiety, which may follow cancer chemotherapy. Cancer-related cognitive impairment (CRCI) has been observed across various cancer therapies, including in individuals with non-central nervous system cancers. Agents such as methotrexate, 5-fluorouracil, and cyclophosphamide have been identified as major contributors to CRCI [5, 6, 12]. This condition is also commonly referred to as “chemobrain” or “chemofog” [4]. Recent studies have reported a high prevalence of chemotherapy-induced cognitive deficits, particularly among breast cancer survivors [13], as well as survivors of other cancer types.

Over the past few decades, research has increasingly focused on the potential of nutraceuticals and dietary supplements to protect against or mitigate the adverse effects of cancer chemotherapy, including hepatotoxicity, subfertility, alopecia, cardiotoxicity, and behavioral deficits [8–11, 14–16].

Dietary supplements such as cholecalciferol, a fat-soluble form of vitamin D, function not only as essential nutrients but also as neurosteroids and hormones [17, 18]. Over the past two decades, numerous studies have demonstrated that vitamin D supplementation enhances both the structure and function of the central nervous system, including improvements in learning and memory [19–24]. The cognitive-enhancing effects of cholecalciferol have been attributed to its ability to cross the blood–brain barrier [22, 23], where it is converted to its active metabolite, 25-hydroxyvitamin D□(25-OH-D□). This metabolite binds to vitamin D receptors, which are widely distributed throughout the brain [18, 24, 25]. Although the neuroprotective properties of cholecalciferol have been reported in several studies using various animal models [18, 26, 27], there remains a significant gap in the literature regarding its potential to ameliorate behavioural deficits induced by cancer chemotherapy. This study, therefore, aimed to evaluate the possible role of vitamin D□in mitigating chemotherapy-associated behavioural impairments in rodents.

## 2. Materials and Methodology

### 2.1 Chemicals and Drugs

Vitamin D3 tablets (1000IU) (Nature’s field®), Cyclophosphamide injection (Endoxan Asta®), Normal saline.

### 2.2 Animals and Animal Welfare

Healthy Wistar rats used in this study were obtained from Empire breeders, Orioke-Ara, Osun State, Nigeria. Rats were housed in wooden cages measuring 20 x 10 x 12 inches in temperature-controlled quarters. All rats were allowed free access to food and water. All procedures were conducted following the approved protocols of the Faculty of Basic Medical Sciences, Ladoke Akintola University of Technology with and within the provisions for animal care and use as prescribed in the scientific procedures on living animals, European Council Directive (EU2010/63).

### 2.3 Experimental Methods

Sixty Wistar rats weighing 180-200 g each were used for the study. The rats were randomly assigned into six groups (n=10) each. Group A (Control Group) received a single intraperitoneal dose of normal saline on the first day and distilled water orally for 15 days. Animals in group B and C received vitamin D_3_ at 300 and 600 IU/kg/day [23, 28]. Group D (cyclophosphamide control) received a single intraperitoneal injection of 100 mg/kg cyclophosphamide on day 1 [23, 29]. Rats in group E and F were administered cyclophosphamide (single dose) with Vitamin D_3_ either at 300 or 600 IU/kg/day received respectively. Intraperitoneal injection of saline or cyclophosphamide was administered as one single dose while, distilled water or Vitamin D_3_ was administered daily for 15 days. At the end of the experimental period, animals were exposed to the behavioural paradigms including the open field arena, Y- and elevated plus mazes.

### 2.4 Behavioural Test

#### 2.4.1 Novelty-induced Open Field Behaviours

Open-field responses in rats are used to depict arousal, inhibitory, diversive, inspective exploratory and anxiety behaviours. Also, stereotypical behaviours like grooming have also been represented using this paradigm. These behaviours are generally regarded as central behaviours and are used to demonstrate the rodent’s ability to explore. Ten minutes of open field behaviours consisting of horizontal locomotion, rearing and grooming, behaviours were monitored and scored in the open field arena as previously described [30, 31]. The open-field paradigm is a square box with a hard floor (covered with glass) that measured 36 x 36 x 26 cm. Wood used was painted white, and its floor was divided by permanent blue markings into 16 equal squares. Each rat was placed in the centre of the field. Total horizontal locomotion (number of floor units entered with all paws), rearing frequency (number of times the rat stood on its hind limbs either with its fore arms against the walls of the observation cage or free in the air) and frequency of grooming (number of body-cleaning with paws, licking of the body and pubis with the mouth, and face-washing actions indicative of a stereotypic behaviour) within the 10 minutes were recorded.

#### 2.4.2 Anxiety Related Behaviours

The elevated plus maze is a plus-shaped apparatus with four arms at right angles to one another. The two open arms lie across from each other, measuring 25×5×5 cm and perpendicular to two closed arms measuring 25×5×16 cm with a central platform (5×5×0.5 cm). The closed arms have a high wall (16 cm) to enclose the arms, whereas the open arms have no side wall. Following administration of drug or vehicle, rats were placed in the central platform facing the closed arm and their behaviour recorded for 5 minutes, as previously described [32, 33]. The criterion for arm visit was considered only when the animal decisively moved all its four limbs into an arm. The maze was cleaned with 5% ethanol following each trial. The percentage of time spent in the arms was calculated as time in open arms or closed arms/total time x100 [31].

#### 2.4.3 Spatial Memory test (Y-maze)

Spatial working memory was assessed using the Y-maze paradigm which is composed of three equally spaced arms (120°, 41cm long and 15cm high). The floor of each arm is made of Perspex and is 5cm wide. Spatial working memory is measured by monitoring spontaneous alternation behaviours. Spatial working memory measured the propensity of rodents to alternate conventionally non-reinforced choices of the Y-maze on successive opportunities. Each rat was placed in one of the arm compartments of the Y maze and allowed to freely move until its tail completely enters another arm. The sequence of arm entries is manually recorded, the arms being labelled A, B, or C. An alternation is defined as entry into all three arms consecutively. The calculation and scoring of the Y maze is as previously reported [34, 35]. For each animal, the Y-maze testing was carried out for 5 minutes. The apparatus was cleaned with 5% alcohol and allowed to dry between sessions.

### 2.5. Statistical Analysis

Data were analysed with Chris Rorden’s ANOVA for Windows (version 0.98). Data analysis was by one-way analysis of variance (ANOVA) and a post-hoc test (Tukey HSD) was used for within and between group comparisons. Results were expressed as mean ± S.E.M. and p < 0.05 was taken as the accepted level of significant difference from control.

## 3 Results

### 4.1 Effect of Cholecalciferol on Behaviours in the Open-Field Paradigm

**Figures 1, 2 and 3** show the effect of cholecalciferol supplementation on open field horizontal locomotion (measured as line crossing), rearing (measured as number of rears), and self-grooming behaviours in CYP-treated rats. Horizontal locomotion increased significantly (p<0.05) with VD at 300 and 600 and decreased with CYP, CYP/VD300, and CYP/VD600 compared to control. Compared to CYP, horizontal locomotion increased with CYP/VD300 and CYP/VD600 respectively. Rearing (vertical locomotion) increased significantly (p<0.05) with VD300 and VD600 and decreased with CYP, CYP/VD300, and CYP/VD600 compared to control. Compared to the CYP, rearing activity increased with CYP/VD600. Self-grooming increased significantly with VD300, VD600, CYP and CYP/VD300 compared to control. Compared to CYP, self-grooming decreased with CYP/VD600.

**Figure 1:**
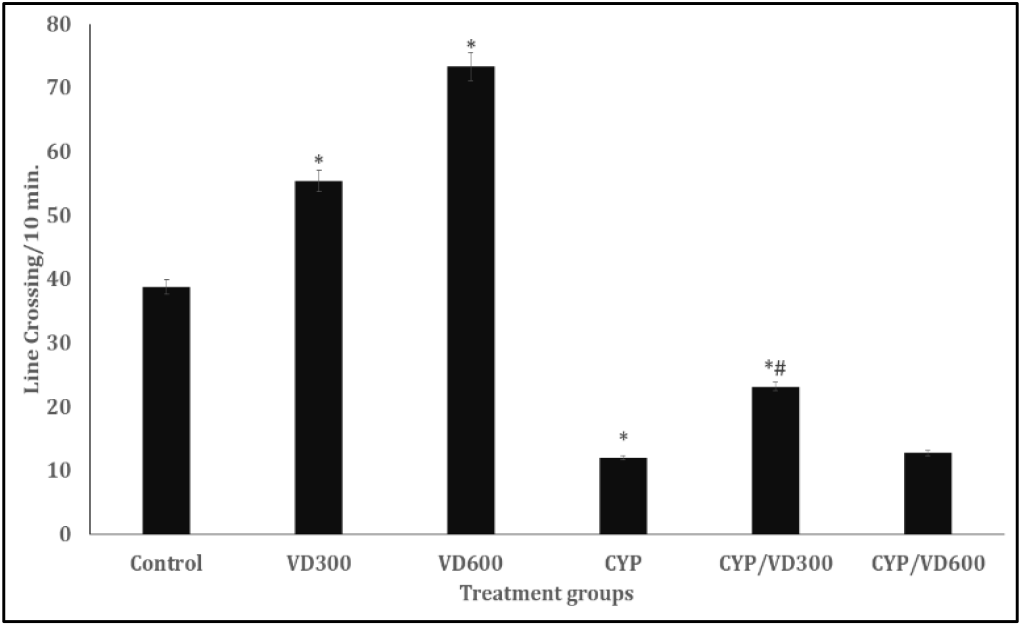
Effect of cholecalciferol on horizontal locomotion measured as line crossing in CYP treated rats. Each bar represents Mean ± S.E.M, * p < 0.05 vs. control, #p<0.05 significant difference from CYP, number of rats per treatment group =10. CONT: Control; CYP: Cyclophosphamide, VD: Vitamin D_3_.

**Figure 2:**
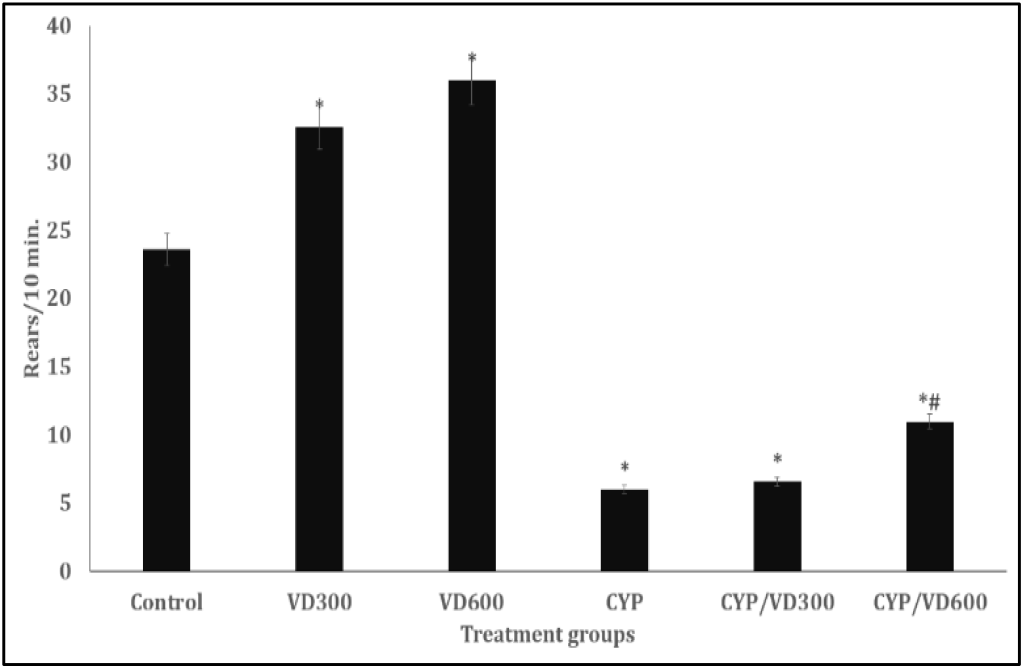
Effect of cholecalciferol on rearing (vertical locomotion). Each bar represents Mean ± S.E.M, * p < 0.05 vs. control, #p<0.05 significant difference from CYP, number of rats per treatment group =10. CONT: Control; CYP: Cyclophosphamide, VD: Vitamin D_3_

**Figure 3:**
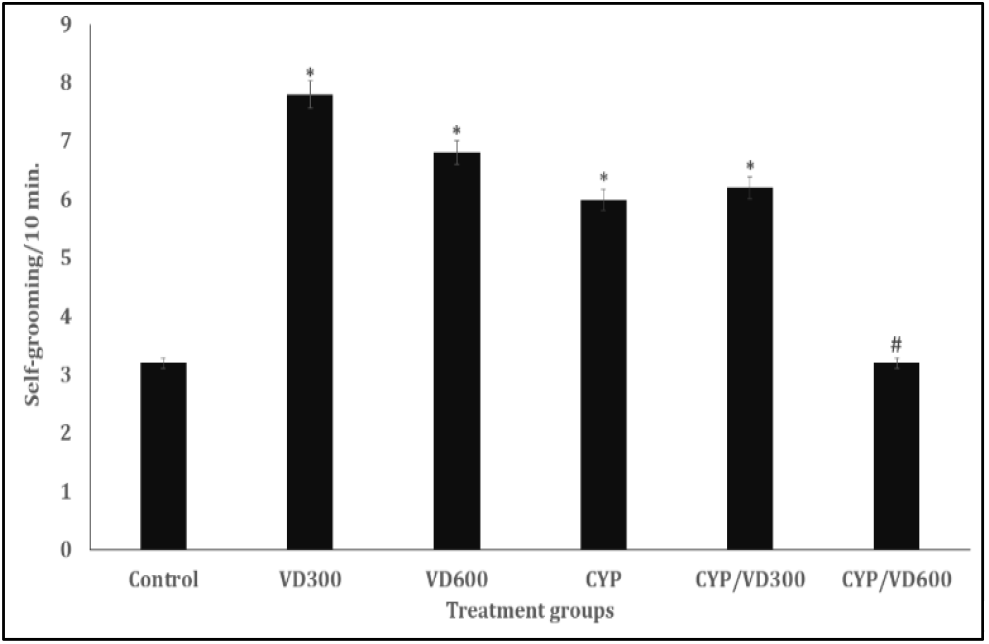
Effect of cholecalciferol on self-grooming. Each bar represents Mean ± S.E.M, * p < 0.05 vs. control, #p<0.05 significant difference from CYP, number of rats per treatment group =10. CONT: Control; CYP: Cyclophosphamide, VD: Vitamin D_3_

### 4.2 Effect of Cholecalciferol on Anxiety-related Behaviours

**Figure 4** shows the effects of cholecalciferol supplementation on anxiety related behaviours measured as percentage time spent in the open arm or closed arm time of the elevated plus maze. Open arm time decreased significantly with VD300, CYP and CYP/VD300 and increased with VD600 and CYP/VD600 compared to control. Compared to CYP, open arm time increased with CYP/VD300 and CYP/VD600. Closed arm time increased with VD300, CYP, and CYP/VD300 and decreased with VD600 and CYP/VD600 compared to control. Compared to CYP, there was a decrease in closed arm time with CYP/VD300 and CYP/VD600 respectively.

**Figure 4:**
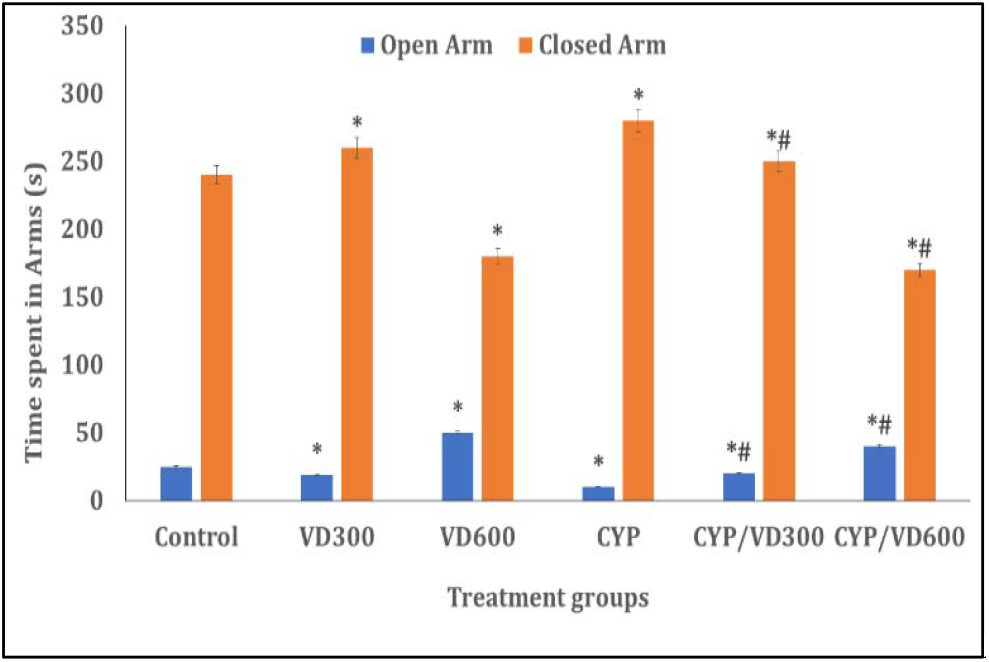
Effect of cholecalciferol on open-arm and closed time in the elevated plus maze. Each bar represents Mean ± S.E.M, * p < 0.05 vs. control, #p<0.05 significant difference from CYP, number of rats per treatment group =10. CONT: Control; CYP: Cyclophosphamide, VD: Vitamin D_3_

### 3.3 Effect of Cholecalciferol on Working Memory

**Figure 5** shows the effect of cholecalciferol supplementation on spatial working memory measured as percentage alternation in the Y-maze. Spatial working memory increased significantly (p<0.05) with VD600, CYP/VD300, CYP/VD600 and decreased with VD300 and CYP compared to control. Compared to CYP, working memory increased with CYP/VD300 and CYP/VD600 respectively.

**Figure 5:**
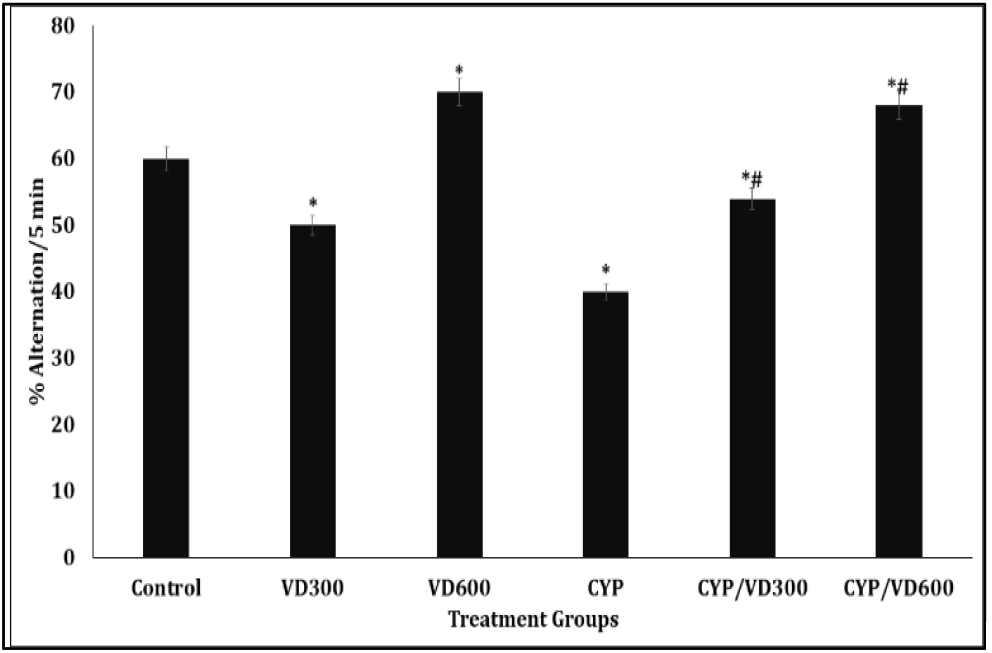
Effect of cholecalciferol on spatial working memory in the Y-maze. Each bar represents Mean ± S.E.M, * p < 0.05 vs. control, #p<0.05 significant difference from CYP, number of rats per treatment group =10. CONT: Control; CYP: Cyclophosphamide, VD: Vitamin D_3._

## 4 Discussion

This study evaluated the neuroprotective effects of cholecalciferol on behavioural toxicities induced by cyclophosphamide treatment in Wistar rats. Cyclophosphamide, a widely used chemotherapeutic agent, has been associated with adverse neurological outcomes, including impairments in learning and memory, reduced motor coordination, and alterations in brain neurochemistry. In the present study, cyclophosphamide administration led to significant behavioural impairments, including decreased horizontal locomotion, reduced rearing activity, memory deficits, and increased anxiety-like behaviours. These findings are consistent with previous studies that have reported the deleterious neurological effects of cyclophosphamide in rodent models [8, 23, 36–38].

Cholecalciferol, an over-the-counter supplement primarily used to correct vitamin D deficiency, has also been shown to exhibit neuroprotective properties [39]. In this study, cholecalciferol administration alone enhanced locomotor activity, rearing, grooming behaviour, and spatial working memory, while also reducing anxiety-related behaviours. These findings align with earlier reports on the cognition-enhancing and anxiolytic effects of cholecalciferol in disease models [19]. Notably, co-administration of cholecalciferol with cyclophosphamide reversed the cyclophosphamide-induced decline in open field activity. Specifically, cholecalciferol mitigated the reduction in line-crossing and rearing behaviours observed in cyclophosphamide-treated rats. An increase in horizontal locomotion and rearing typically reflects central excitatory responses, whereas a decrease suggests central inhibitory effects. These behavioural outcomes are indicative of underlying neural mechanisms that modulate postsynaptic activity either through excitation or inhibition [40]. The hypolocomotor effects of cyclophosphamide observed in this study have also been reported in other preclinical studies [8, 41]. Furthermore, cholecalciferol effectively reversed the cyclophosphamide-induced impairments in spatial working memory and anxiety-related behaviours, as assessed using the Y-maze and behavioural anxiety indices. Previous studies have similarly linked cyclophosphamide treatment with cognitive deficits and heightened anxiety [42, 43]. The ability of cholecalciferol to mitigate these behavioural abnormalities is likely mediated by its antioxidant and anti-inflammatory properties, which help counteract cyclophosphamide-induced oxidative stress and neuroinflammation [44].

## Conclusion

In conclusion the results of this study revealed that cholecalciferol therapy has the ability to mitigate cyclophosphamide-induced behavioural deficits. However, more studies would be needed to ascertain its benefits in humans.

## Acknowledgements

**None**

## Availability of data and materials

Data generated during and analysed during the course of this study are available from the corresponding author on request.

## Declaration of ethical approval

Ethical approval for this study was granted by the Ethical Committee of the Faculty of Basic Medical Sciences ERC/FBMS/038/2024

## Competing interests

None

Received: 7^th^ May 2025, revised: 5^th^ June 2025, Accepted:28^th^ June, 2025. Published online

